# Targeting the TRA-1-60 Glycoepitope Enables Selective ImmunoPET Imaging of Ovarian Cancer

**DOI:** 10.64898/2026.08.12.744522

**Authors:** Sajmina Khatun, Alexandra Fox, Alexandra Skowron, Ayesha B. Alvero, Nerissa Viola

## Abstract

Targeted radiopharmaceutical development for ovarian cancer (OC) has been limited by the lack of molecular targets that combine broad tumor expression with minimal normal-tissue distribution. TRA-1-60 (TRA) is a cancer-associated glycoepitope carried by podocalyxin. Here, we evaluated TRA as a target for OC and developed a TRA-directed immunoPET imaging platform. Immunohistochemical analysis demonstrated significantly higher TRA expression in ovarian tumors than in normal adjacent ovarian tissue, with expression maintained across epithelial OC histotypes and disease stages. An engineered anti-TRA single-chain variable fragment-Fc (scFv-Fc) demonstrated robust penetration of three-dimensional tumor spheroids and selective accumulation in intraperitoneal tumors in an immunocompetent syngeneic OC model. Radiolabeling with zirconium-89 generated [⁸⁹Zr]Zr-DFO-anti-TRA scFv-Fc with >98% radiochemical yield. Serial PET/CT imaging demonstrated progressive and sustained radiotracer accumulation at tumor sites through 96 hours, accompanied by declining liver-associated activity and low uptake in most normal tissues. Together, these findings identify TRA as a broadly expressed and accessible tumor-associated glycoepitope and establish TRA-targeted immunoPET as a promising strategy for noninvasive detection of OC. The selective and sustained tumor localization of this platform further provides a foundation for development of TRA-directed radiopharmaceutical therapy, supporting a potential theranostic approach for OC.

## Introduction

Recurrent ovarian cancer (OC) remains a major clinical challenge and a leading cause of gynecologic cancer mortality ^1–3^. Despite initial responses to platinum-based chemotherapy, most patients ultimately relapse and develop chemoresistant disease, at which point treatment options are limited and clinical outcomes remain poor ^4,5^. The incorporation of targeted therapies such as bevacizumab and poly-ADP ribose polymerase (PARP) inhibitors has resulted in only modest improvements in overall survival ^5,6^. More recently, immune checkpoint blockade with the PD-1 inhibitor pembrolizumab (Keytruda®) has received FDA approval in select settings; however, the overall clinical benefit of immunotherapy in OC has been limited by low response rates, intrinsic immune resistance, and restricted patient eligibility ^7^. Thus, there remains a critical unmet need for more effective and broadly applicable therapeutic strategies for recurrent and platinum-resistant OC.

Targeted radiopharmaceutical agents are an emerging precision oncology platform that combine molecular targeting with radioactive payloads to enable tumor-specific imaging and therapy ^8,9^. These agents consist of a targeting moiety linked to a radionuclide that can function either as a diagnostic imaging tracer or as a therapeutic agent delivering localized DNA-damaging radiation directly to tumor cells. The clinical success of targeted radiopharmaceuticals in other malignancies has established this modality as a transformative approach in cancer management ^9^. In prostate cancer, PSMA-targeted agents such as [⁶⁸Ga]Ga-PSMA-11 and [¹⁸F]DCFPyL are FDA-approved for positron emission tomography (PET) imaging, while [¹⁷⁷Lu]Lu-PSMA-617 (Pluvicto®) is approved for metastatic castration-resistant disease ^10^. Similarly, in neuroendocrine tumors, somatostatin receptor-targeted agents including [⁶⁸Ga]Ga-DOTATATE and [¹⁷⁷Lu]Lu-DOTATATE (Lutathera®) have demonstrated substantial clinical benefit ^11^. Together, these examples highlight the ability of targeted radiopharmaceuticals to deliver highly selective systemic therapies while minimizing off-target toxicity.

In contrast, targeted radiopharmaceutical agents remain relatively underexplored in OC ^12^. Although several OC-directed radiopharmaceutical therapies have shown promise in preclinical studies, only a limited number have advanced into early-phase clinical trials. Radioligands targeting HER2 or NaPi2b have demonstrated favorable therapeutic windows in Phase I studies for ovarian and peritoneal malignancies ^13,14^. Nevertheless, broader clinical translation of radiopharmaceuticals in OC has been hindered by the lack of tumor-selective, clinically actionable targets. This challenge is compounded by the profound molecular heterogeneity of OC and its characteristic diffuse intraperitoneal (i.p) dissemination ^15–17^, both of which necessitate targets that provide broad tumor coverage while maintaining high specificity.

Current OC-targeted therapies further illustrate this challenge. Folate receptor alpha (FRα), the target of the FDA-approved antibody-drug conjugate Elahere®, has validated the clinical utility of tumor-directed therapy in OC. However, its heterogeneous expression across tumors and presence in normal tissues, including renal tubular epithelium and the retina ^18^ highlight potential limitations related to patient coverage and off-tumor effects. Thus, identifying targets that combine broad tumor expression with minimal normal-tissue distribution remains a critical priority for developing safer and more broadly applicable targeted therapies for OC.

To address this unmet need, we developed a targeted radiopharmaceutical strategy directed against TRA-1-60 (TRA), a cancer-associated glycoepitope expressed on the heavily glycosylated cell-surface protein podocalyxin (PODXL). Unlike PODXL, which is expressed in both normal and malignant tissues ^19,20^, the TRA glycoepitope is minimally expressed in healthy tissues, and heavily expressed in cancer cells reflecting the cancer-associated aberrant glycosylation that accompanies malignant transformation ^21,22^. In this study, we demonstrate that the TRA glycoepitope is broadly expressed across OC histotypes, and exhibits highly selective tumor localization *in vivo* with minimal expression in normal ovarian tissues. ImmunoPET imaging with a ^89^Zr-labeled anti-TRA single-chain variable fragment-Fc (scFv-Fc) targeting radioligand enabled specific and sustained visualization of i.p. ovarian tumors with favorable pharmacokinetic properties.

## Materials and Methods

### Ovarian cancer tissue microarray (TMA) and immunohistochemistry (IHC)

The OC TMA (OV1006) was purchased from TissueArray.com and immunostained with TRA antibody (RRID:AB_891610) at a 1:50 dilution. Heat-induced epitope retrieval was performed at 95°C using pH ∼9 retrieval buffer. Staining was developed using the Novolink Polymer Detection System (Leica Biosystems, RE7280-CE). Slides were scanned using the Aperio Scanner CS2 (RRID:SCR_025111), and TRA staining was quantified using HALO (RRID:SCR_018350), with the CytoNuclear v2.0.9 module (Indica Labs).

### Ovarian cancer cell lines and culture methods

Human OC cell lines SKOV3 (RRID:CVCL_0532), OVCAR3 (RRID:CVCL_0465), OVCA432 (RRID:CVCL_3769), PE01 (RRID:CVCL_2686), PE04 (RRID:CVCL_2690) and R182 (RRID: CVCL_6B94)^23–27^ were grown in Roswell Park Memorial Institute (RPMI) 1640 media containing 10% fetal bovine serum (FBS), 1% penicillin-streptomycin, 1% MEM-NEAA, 1% HEPES and 1% sodium pyruvate. Triple-knockout (TKO) mouse OC cells were kindly provided by Dr. M. Matzuk ^28^. TKO cells were maintained in a 1:1 mixture of Dulbecco’s Modified Eagle Medium (DMEM) and Ham’s F12 medium supplemented with 10% FBS and 1% penicillin–streptomycin. Stable expression of mCherry fluorescence in TKO cells was achieved using lentiviral transduction as previously described ^29^. All cell lines were maintained at 37°C in a humidified incubator with 5% CO₂. Cultures were tested for Mycoplasma contamination monthly and authenticated annually by short tandem repeat (STR) profiling. Cells were used within eight passages for all experiments to ensure phenotypic consistency.

### Protein extraction and quantification

Whole-cell protein lysates were prepared using 10× Cell Lysis Buffer (Cell Signaling Technology, Danvers, MA) according to the manufacturer’s instructions. Briefly, cell pellets were resuspended in ice-cold 1× lysis buffer supplemented with Complete™ Protease Inhibitor Cocktail (Millipore Sigma, Burlington, MA), incubated on ice, and centrifuged at 13,000 rpm for 15 minutes at 4°C to remove cellular debris. Supernatants containing soluble proteins were collected, and protein concentrations were determined using the Pierce BCA Protein Assay Kit (Thermo Fisher Scientific, Waltham, MA).

### SDS-PAGE and western blot analysis

Equal amounts of protein lysates (20 μg) were separated by electrophoresis on 10% SDS-polyacrylamide gels and transferred onto PVDF membranes (EMD Millipore, Burlington, MA). Membranes were blocked in 5% non-fat milk in phosphate-buffered saline (PBS)-Tween then incubated overnight at 4°C with primary antibody against human PODXL (RRID: AB_354920) orβ-actin (RRID: AB_10700003). Following washing, membranes were incubated with appropriate HRP-conjugated secondary antibodies for 1 hour at room temperature. Protein bands were detected using enhanced chemiluminescence reagent and visualized with the ImageQuant LAS 500 (Cytiva Life Sciences, Marlborough, MA).

### Generation of anti-TRA scFv-Fc antibody and conjugation with Cy5 or Cy7

Bstrongomab (Bsg; Curemeta, LLC) is a humanized IgG1 antibody directed against the TRA epitope. The anti-TRA scFv-Fc fragment (110 kDa) derived from the Bsg antibody was conjugated to Cy5 or Cy7 at a mole ratio of 1:10 by incubating at 37°C with gentle shaking at 300 rpm for 1 hour. Following incubation, the conjugated product was purified using Zeba Spin columns according to the manufacturer’s purification protocol with saline exchange.

### 3D spheroids and immunofluorescence

Three-dimensional cancer spheroids were generated by seeding 2,500 cells per well into 96-well round-bottom ultra-low attachment plates (Corning Life Sciences, Glendale, AZ) and cultured for 96 h to allow spheroid formation. Spheroids were then transferred to 24-well plates containing complete growth medium and incubated with Cy5-conjugated anti-TRA scFv-Fc for 1 h under standard culture conditions. After washing with 1× PBS to remove unbound antibody, fluorescence images were acquired to assess antibody binding and penetration throughout the spheroids. All experiments were performed in three independent biological replicates.

### Establishment of mouse models of OC

All animal studies were conducted in accordance with protocols approved by the Wayne State University Institutional Animal Care and Use Committee (IACUC; Protocols 25-01-7461 and 23-10-6206). Mice were maintained under standard housing conditions with *ad libitum* access to food and water and were acclimated for at least one week prior to tumor implantation.

For the xenograft model, 4 × 10⁶ SKOV3 cells were resuspended in a 1:1 mixture of growth medium and Geltrex™ basement membrane matrix (Thermo Fisher Scientific) and injected subcutaneously (s.c.) into the right shoulder of 6-week-old female athymic nude mice (RRID: IMSR_ENV:HSD-069). Tumor growth was monitored twice weekly by palpation and caliper measurements. Once tumors reached a volume of 80–150 mm³, they were excised and processed for IHC analysis.

For the syngeneic tumor model ^30–33^, 1 × 10⁷ mCherry-expressing TKO mouse OC cells were injected i.p. into 8-week-old female C57BL/6 mice (RRID: IMSR_JAX:000664). Tumor progression was monitored twice weekly by *in vivo* fluorescence imaging (570-nm excitation/630-nm emission) under isoflurane anesthesia using the Ami HT Imaging System (Spectral Instruments Imaging) and mCherry signal was quantified as regions of interest (ROI) using Aura Imaging Software. All mice meeting predefined tumor establishment criteria (3.4 × 10⁸ photons/sec mCherry ROI) were included in the study and randomized into groups. Investigators were not blinded to group allocation. Animals were monitored closely throughout the study for signs of distress, and humane endpoints were implemented in accordance with IACUC guidelines.

### Administration of fluorescent probe and ex vivo biodistribution analysis

Mice bearing TKO tumors were administered a single i.v. dose of 200 μg Cy7-conjugated anti-TRA scFv-Fc. Ninety-six hours following antibody administration, mice were euthanized and subjected to necropsy for *ex vivo* biodistribution analysis. Major organs and omental metastasis were dissected and imaged to detect both mCherry fluorescence, indicative of tumor burden, and Cy7 fluorescence, corresponding to antibody distribution. Fluorescence images were acquired under standardized exposure settings, and ROI were drawn for each tissue to quantify fluorescence intensity using Aura Imaging Software. ROI measurements were used to assess tissue biodistribution and tumor-specific accumulation of the Cy7-conjugated antibody.

### Conjugation of DFO Chelator to the anti-TRA scFv-Fc Antibody

The anti-TRA scFv-Fc was functionalized with the bifunctional chelator p-isothiocyanatobenzyl-desferrioxamine (DFO-Bz-SCN; Macrocyclics) following previously established protocols^34^. Conjugation was performed by dissolving the antibody in 0.9% (w/v) sterile saline adjusted to pH ∼9, followed by the addition of DFO dissolved in dimethyl sulfoxide (DMSO) at an antibody-to-chelator molar ratio of 1:5. The reaction mixture was incubated at 37 °C for 1 h under gentle agitation to facilitate isothiocyanate-mediated covalent coupling to the primary amines of the antibody. Upon completion of the incubation, unbound and excess DFO was removed by centrifugal ultrafiltration using a membrane with a molecular weight cut-off (MWCO) of 30 kDa, with 0.9% sterile saline employed as the eluent. The resulting immunoconjugate, designated as DFO-anti-TRA scFv-Fc was collected and stored at 4 °C until further use.

### ⁸⁹Zr Radiolabeling

Radiolabeling of the DFO-anti-TRA scFv-Fc with zirconium-89 was performed under neutral aqueous conditions. Briefly, approximately 1 mCi (37 MBq) of [⁸⁹Zr]Zr-oxalate was neutralized to a pH of 7.0-7.2 by the dropwise addition of 1 M sodium carbonate . The pH-adjusted [⁸⁹Zr]Zr solution was subsequently combined with DFO-anti-TRA scFv-Fc (0.2 mg, 2 nmol) and incubated at room temperature for 1 h to allow for efficient chelation of ⁸⁹Zr onto the DFO chelate. The crude radiolabeled product, [⁸⁹Zr]Zr-DFO-anti-TRA scFv-Fc, was purified by centrifugal spin-column filtration using a 7 kDa MWCO membrane, with sterile 0.9% saline as the eluent, to remove low-molecular-weight impurities and unincorporated radiometal. The radiochemical purity (RCP) of the final radiolabeled construct was determined by radio-instant thin-layer chromatography (radio-iTLC) using a silica-gel impregnated glass microfiber chromatography paper strip (Agilent iTLC-SG) developed using 50 mM EDTA as a mobile phase.

### PET/CT Imaging and tissue biodistribution

Mice bearing TKO tumors received an intravenous injection of [⁸⁹Zr]Zr-DFO-anti-TRA scFv-Fc-(200 μCi per mouse) in sterile saline. Longitudinal small-animal PET imaging was performed at 3, 24, 48, 72, and 96 h post-injection (p.i.) to evaluate the *in vivo* biodistribution, tumor uptake, and retention of the radiotracer over time. PET images were acquired using a Bruker Albira Si PET/CT scanner with mice that are anesthetized via inhalation of 2% isoflurane in O_2_ throughout the imaging acquisition to minimize motion artifacts. PET data were reconstructed using the Maximum Likelihood Expectation Maximization (MLEM) algorithm (12 iterations; 0.75-mm voxel resolution). Reconstructed images were analyzed with Imalytics Preclinical software (RRID:SCR_026980)by manually delineating volumes of interest (VOIs) over tumors and selected organs on serial image slices and expressed as the percentage of injected dose per volume of tissue (% ID/mL), enabling quantitative comparison of tumor uptake and biodistribution across imaging time points.

Tissue biodistribution of [⁸⁹Zr]Zr-DFO-anti-TRA scFv-Fc was evaluated at 24, 48, and 96 h p.i. A 100-fold excess unmodified anti-TRA scFv-Fc was co-administered with the radiotracer in a separate group of mice to test for specificity at 48 h p.i. Following euthanasia via CO_2_ asphyxiation, blood was collected by cardiac puncture while tumors and selected organs were excised, weighed, and analyzed for bound radioactivity using a calibrated gamma counter. Radioactivity measurements were corrected for background and the physical decay of ⁸⁹Zr. Radiotracer uptake was expressed as the percentage of injected dose per gram of tissue (%ID/g), allowing quantitative comparison of tumor targeting and normal tissue biodistribution across time points.

### Statistical analysis

Statistical analyses were performed using GraphPad Prism (RRID:SCR_002798). Data are presented as mean ± standard error of the mean (SEM) unless otherwise indicated. Comparisons between two groups were analyzed using an unpaired two-tailed Student’s *t*-test. Comparisons among three or more groups were performed using one-way analysis of variance (ANOVA) followed by appropriate post hoc multiple-comparison testing. A *p* value < 0.05 was considered statistically significant.

## Results

### TRA differentially recognizes ovarian tumors from normal adjacent ovarian tissue

To evaluate whether TRA is a useful and specific targeting marker for OC, we assessed expression of the TRA-reactive epitope by IHC using a comprehensive OC TMA. The TMA included primary tumors representing all major epithelial OC histologic subtypes across disease stages, as well as metastatic lesions from omentum and lymph node, and normal adjacent ovarian tissue (NAT) (Table 1). TRA exhibited prominent membranous staining in malignant epithelial cells, whereas NAT showed minimal to absent staining (Fig. 1A). Quantification of staining intensity using H-scores demonstrated significantly higher TRA expression in primary tumors compared with NAT (Fig. 1B). This difference remained significant when analysis was restricted to high-grade serous ovarian cancer (HGSOC) (Fig. 1C). TRA expression did not vary significantly between early-stage (Stage I–II) and late-stage (Stage III–IV) HGSOC, nor among the different epithelial OC histologic subtypes (Fig. 1D-E). These findings establish TRA as a broadly expressed, tumor-associated glycoepitope that distinguishes ovarian tumors from NAT and is conserved across OC histotypes and disease stages.

**Figure 1.**
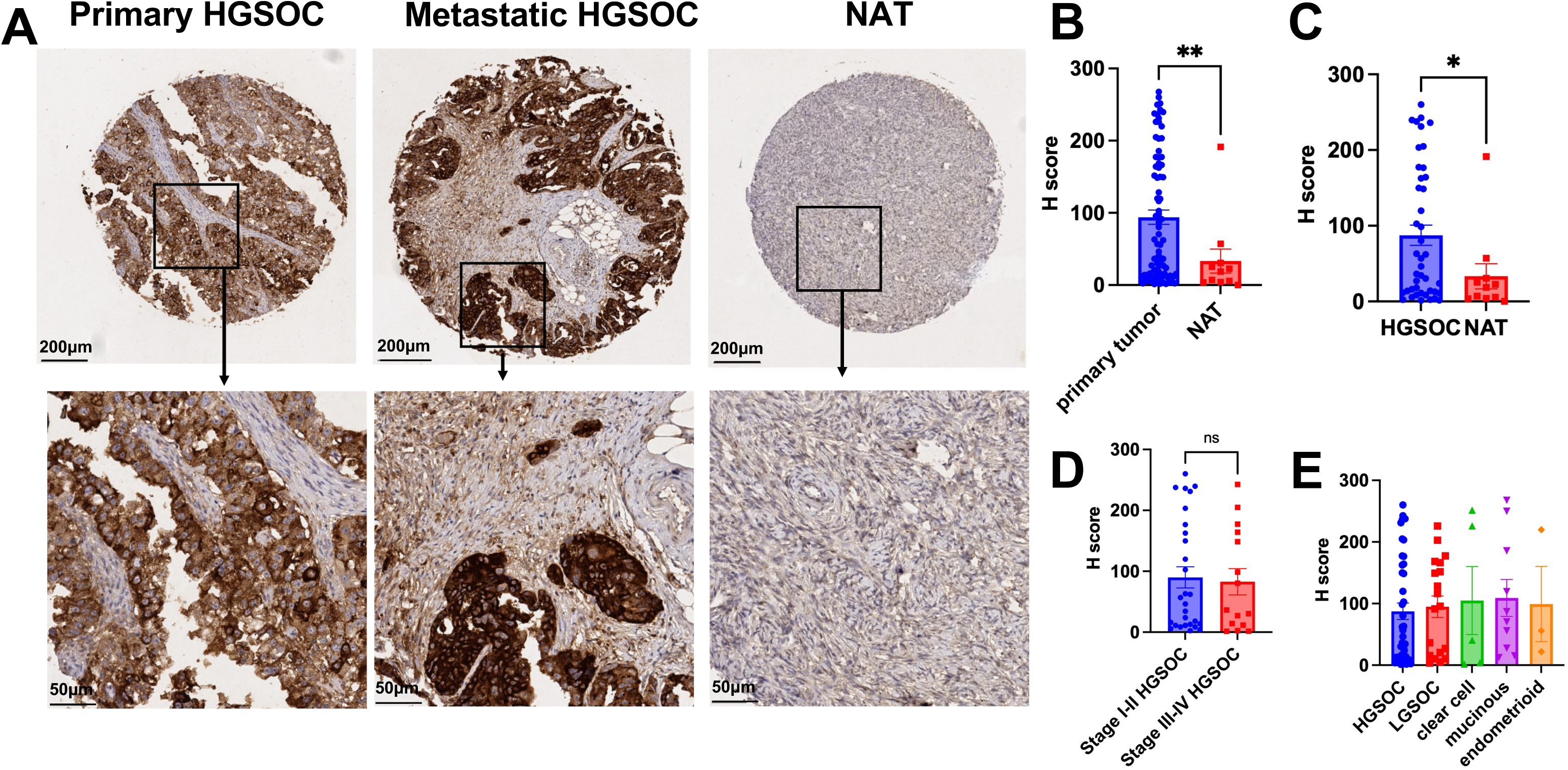
TRA expression distinguishes primary and metastatic ovarian tumors from normal adjacent ovarian tissue. Epithelial ovarian cancer TMA were immunostained with antibody against TRA, and H-scores were quantified. **(A)** Representative images from TMA cores showing intense membrane staining in primary high-grade serous ovarian cancer (HGSOC) and metastatic lesions but not normal adjacent ovarian tissue (NAT); **(B)** TRAH-scores were significantly higher in primary tumors vs NAT; **(C)** This difference remained significant when the analysis was restricted to HGSOC vs NAT; **(D)** No significant differences in TRA H-scores were observed between early-stage (Stage I–II) and late-stage (Stage III–IV) HGSOC, or **(E)** among the various epithelial ovarian cancer histologic subtypes.

**Table 1.** Composition of the Ovarian Cancer Tissue Microarray by Histological Subtype and Disease Stage.

| **Primary tumor** |  | *(n)* |
| --- | --- | --- |
|  | High-grade serous | 43 |
|  | Low-grade serous | 19 |
|  | Endometrioid | 3 |
|  | Mucinous | 10 |
|  | Clear Cell | 5 |
| **Metastasis** |  |  |
|  | Omentum | 5 |
|  | Lymph node | 5 |
| **Norma adjacent ovarian tissue (NAT)** |  | 10 |
| **Stage** | I | 36 |
|  | II | 12 |
|  | III | 18 |
|  | IV | 4 |

### TRA exhibits robust cell-surface accessibility in human SKOV3 OC xenograft model

To further determine the utility of TRA as a targeting agent for OC, we sought to identify relevant preclinical OC models with robust expression of the TRA-reactive epitope. Since TRA is a glycosylated epitope carried by PODXL, we evaluated PODXL expression across a panel of human OC cell lines by western blot analysis. All cell lines examined, with the exception of OVCAR3, showed detectable levels of PODXL, although the apparent molecular weight varied among cell lines (Fig. 2A). The observed differences in electrophoretic mobility on SDS-PAGE likely reflect heterogeneous glycosylation patterns, consistent with the heavily glycosylated nature of PODXL ^35^.

**Figure 2.**
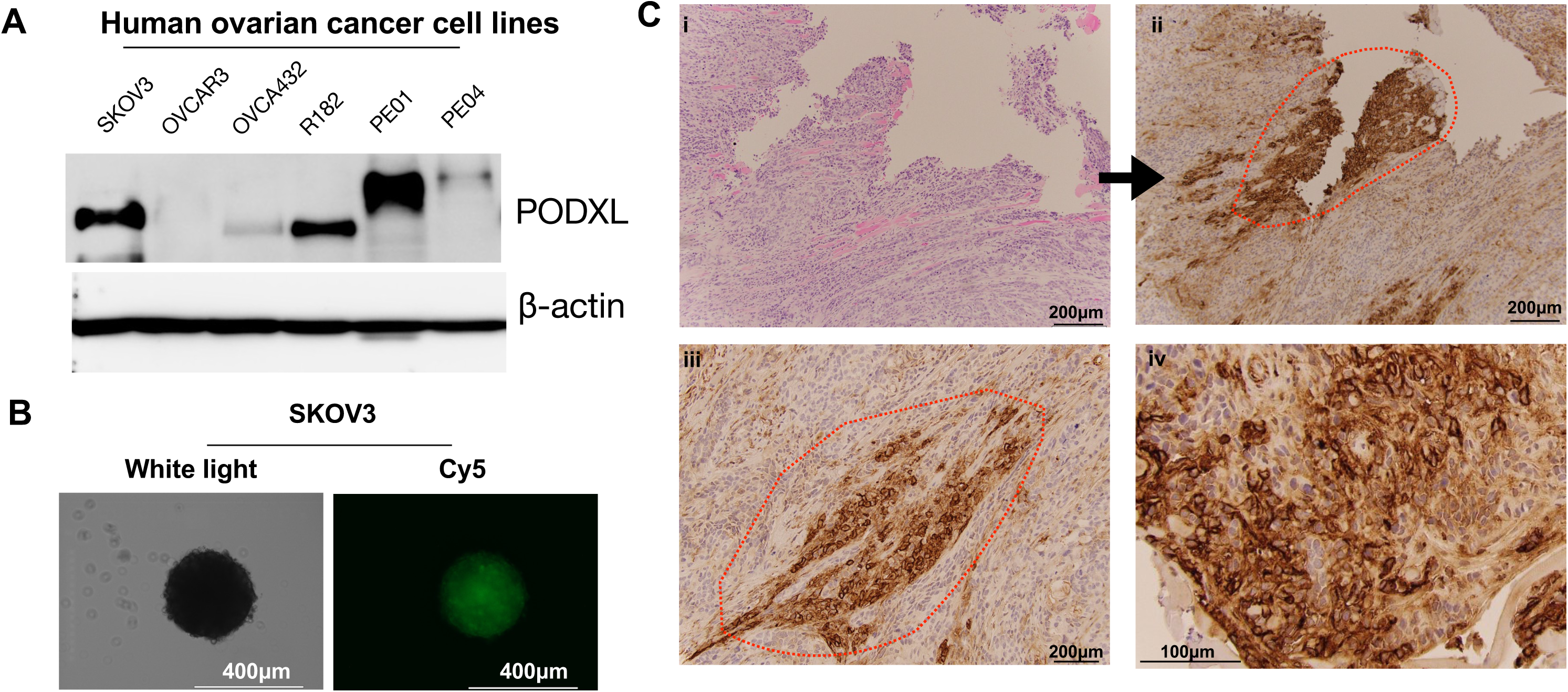
TRA is robustly expressed on human ovarian cancer cells in 3D spheroids, and *in vivo*. **(A)** Expression of PODXL, the carrier protein for the TRA epitope, across a panel of human ovarian cancer cell lines**; (B)** PODXL-positive human SKOV3 OC cells were cultured in ultra-low attachment plates to generate spheroids and subsequently immunostained with Cy5-conjugated anti-TRA scFv-Fc (detailed in text), demonstrating robust surface expression in 3D spheroids**; (C)** SKOV3 OC cells were injected s.c. into athymic nude mice, and resulting tumors were analyzed by H&E (i) and IHC staining with anti-TRA (ii-iv). TRA staining was predominantly localized to the cell membrane, consistent with surface expression of the antigen *in vivo*. Red dashed circle, epithelial tumors.

Among the PODXL-positive cell lines, SKOV3 cells were selected for further studies because they reliably and reproducibly form tumors *in vivo* ^36^. To better model the 3D tumor microenvironment and predict *in vivo* targeting, uptake studies were performed using SKOV3 tumor spheroids. These studies utilized a Cy5-conjugated humanized anti-TRA scFv-Fc . As shown in Figure 2B, the Cy5-conjugated anti-TRA scFv-Fc probe demonstrated robust and uniform uptake throughout intact, well-formed tumor spheroids. Bright and homogeneous fluorescence signals were observed across the entire spheroid and closely corresponded with the compact spheroid morphology observed by brightfield imaging. The widespread distribution of fluorescence within the spheroid suggests efficient probe penetration and retention throughout the 3D structure. This finding is particularly significant given that limited tissue penetration is a major challenge for antibody-based imaging and therapeutic agents.

To determine whether TRA expression is maintained *in vivo*, SKOV3 cells were injected s.c. into athymic nude mice, and the resulting tumors were analyzed by IHC for TRA. Robust TRA staining was observed throughout malignant epithelial tumor regions (Fig. 2C, dashed red line), whereas surrounding stromal areas demonstrated minimal staining. Importantly, TRA expression was predominantly localized to the cell membrane, consistent with the cell-surface distribution required for efficient antibody-mediated targeting. Together with the spheroid uptake studies, these findings identify TRA as a highly accessible tumor-associated surface epitope with favorable characteristics for targeting OC.

### Anti-TRA scFv-Fc selectively localized to tumors in an orthotopic immunocompetent OC model

We next sought to determine whether TRA accessibility was maintained in an orthotopic OC model. For this, we used the TKO mouse OC cells derived from spontaneously arising HGSOC in mice harboring conditional deletions of *Dicer* and *Pten* together with a gain-of-function *Trp53* mutation ^28,37^. TKO spheroids, despite exhibiting a more loosely organized architecture than SKOV3 spheroids, demonstrated similarly robust uptake and penetration of the Cy5-conjugated anti-TRA scFv-Fc (Fig. 3A–B), indicating that efficient TRA targeting is preserved across distinct OC spheroid models.

**Figure 3.**
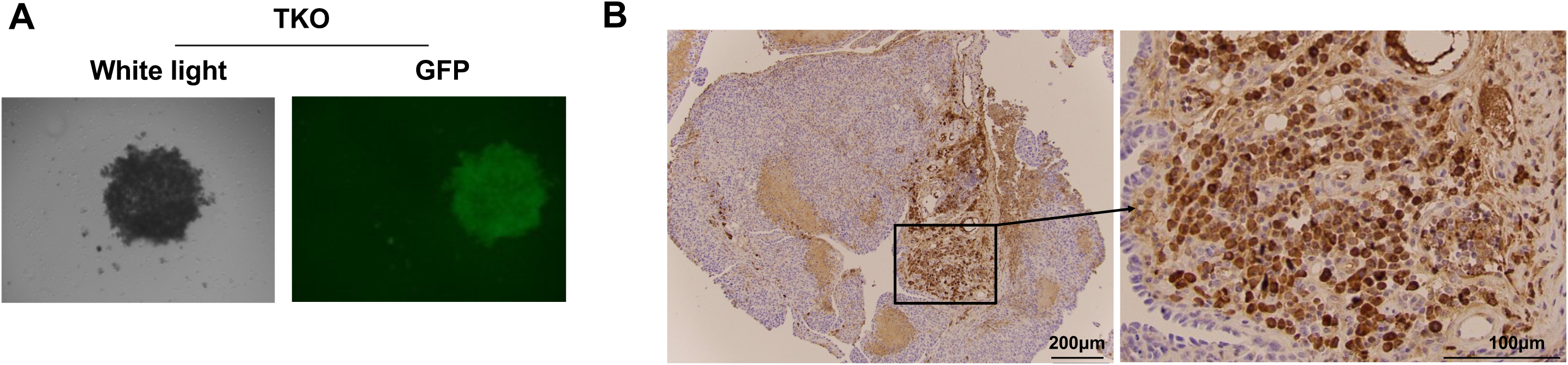
TRA is expressed on murine ovarian cancer spheroids and omental metastases *in vivo*. **(A)** Mouse TKO ovarian cancer cells were cultured in ultra-low attachment plates to generate multicellular spheroids and subsequently immunostained with Cy5-conjugated anti-TRA scFv-Fc **; (B)** TKO OC cells were injected intraperitoneally into C57BL/6 mice to establish omental metastases, which were subsequently analyzed by IHC staining with anti-TRA.

To confirm that TRA expression was retained *in vivo*, i.p. TKO tumors were harvested and analyzed by IHC. Orthotopic TKO tumors demonstrated positive TRA staining within the tumor tissue (Fig. 3B). TRA expression was localized predominantly to malignant epithelial tumor cells, whereas the surrounding stromal tissue exhibited minimal staining. Although staining intensity was heterogeneous across the tumor, distinct regions showed strong positive immunoreactivity, indicating that the TRA epitope is retained *in vivo* and remains accessible on tumor cells within the orthotopic OC microenvironment.

We next evaluated the *in vivo* tumor-targeting capability of the anti-TRA scFv-Fc. Mice bearing established i.p. TKO tumors received a single i.p. injection of Cy7-conjugated anti-TRA scFv-Fc. *Ex vivo* fluorescence imaging performed 96 h later demonstrated selective accumulation of the probe within mCherry-positive tumor implants (Fig. 4A,B). Minimal signal was detected in the liver and kidneys, which are common sites of nonspecific uptake and clearance for antibody-based imaging agents. Quantitative ROI analysis confirmed high tumor-to-liver and tumor-to-kidney fluorescence ratios, indicating preferential uptake and prolonged retention of the probe within ovarian tumors (Fig. 4C,D).

**Figure 4.**
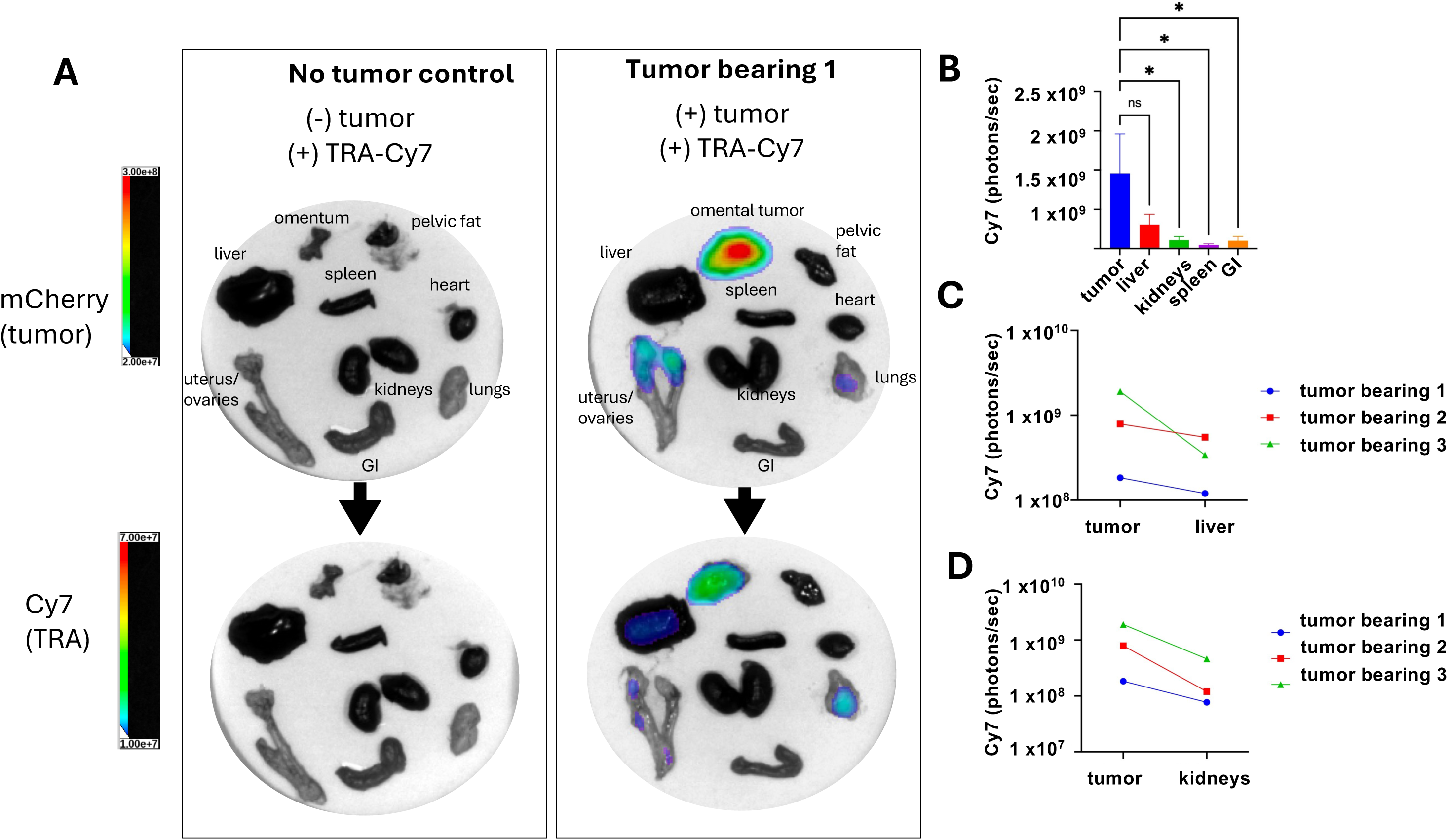
*In vivo* biodistribution of Cy7-conjugated anti-TRA scFv-Fc in an orthotopic OC model. TKO OC cells were injected intraperitoneally into C57BL/6 mice, and tumor growth was monitored longitudinally by *in vivo* imaging of the mCherry signal. Once mice developed measurable disease (mCherry signal ≥ 3.4 × 10^8 photons/sec), animals received a single intraperitoneal dose (100 μg) of Cy7-conjugated anti-TRA scFv-Fc (n=3). Ninety-six hours after antibody administration, mice were euthanized and tissues were collected at necropsy for *ex vivo* biodistribution analysis. Cy 7 fluorescence intensity was quantified to evaluate tissue distribution and tumor-specific accumulation of the antibody. * p<0.05 by One Way ANOVA.

### PET Radiolabeling and PET/CT Imaging

Having established robust and selective tumor accumulation of anti-TRA scFv-Fc in the TKO model, we next evaluated whether this construct could be adapted into a clinically relevant PET imaging radiotracer. Compared with optical imaging, PET enables sensitive, quantitative, whole-body assessment of radiotracer distribution with greater tissue penetration, making it better suited for clinical translation. To generate a TRA-targeted immunoPET tracer, the DFO-conjugated anti-TRA scFv-Fc construct was radiolabeled with zirconium-89 (^89^Zr), a positron-emitting radionuclide well-suited for immunoPET imaging due to its favorable half-life and compatibility with antibody-based tracers. Radiolabeling was robust and reproducible, achieving a radiochemical yield of >98%. The resulting radiotracer, [^89^Zr]Zr-DFO- anti-TRA scFv-Fc, was obtained with a specific activity of 185 ± 10 MBq/mg.

To evaluate tumor targeting *in vivo*, mice bearing established i.p. TKO tumors received a single i.v. injection of 200 μCi [⁸⁹Zr]Zr-DFO-anti-TRA scFv-Fc, followed by serial PET/CT imaging at 3, 24, 48, 72, and 96 h post-injection (p.i.) (n=4). Representative mCherry fluorescence at the time of radiotracer administration confirmed tumor location (Fig. 5A), which for this model is typically the omentum and adjacent spleen ^30,31,33^ . Serial PET/CT imaging demonstrated radiotracer accumulation at the corresponding tumor site that persisted throughout the 96-h imaging period (Fig. 5B, dashed circles). Quantitative volume-of-interest (VOI) analysis confirmed sustained tumor-associated signal, with radiotracer uptake increasing at later time points (Fig. 5C). In contrast, liver-associated signal progressively declined over time (Fig. 5B, D), resulting in improved tumor-to-liver contrast. Together, these findings demonstrate progressive and sustained accumulation of [⁸⁹Zr]Zr-DFO-anti-TRA scFv-Fc in i.p. TKO tumors, accompanied by clearance from the liver, supporting favorable *in vivo* pharmacokinetics and selective targeting of TRA-expressing ovarian tumors.

**Figure 5.**
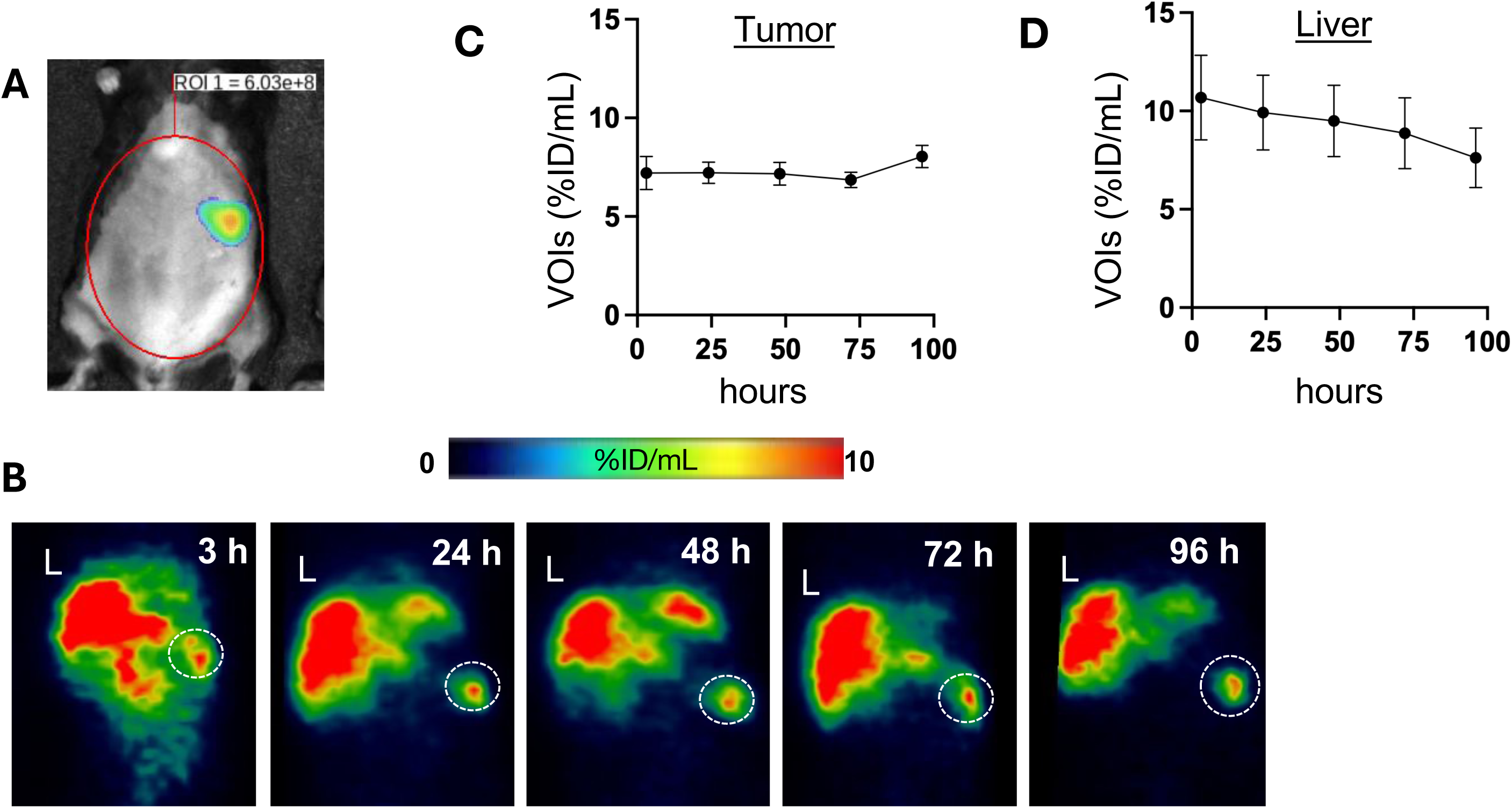
TRA-targeted radiotracer ([^89^Zr]Zr-DFO-anti-TRA scFv-Fc) demonstrates progressive tumor accumulation and clearance from the liver. Mice bearing i.p. TKO tumors received a single i.p. dose (300 μCi) of [^89^Zr]Zr-DFO-anti-TRA scFv-Fc (n=3). **(A)** Representative mCherry fluorescence imaging at the time of radiotracer administration identifies tumor localization, primarily within the omentum, a characteristic site of tumor implantation in this model ^30,31,33^; **(B)** Representative serial PET/CT images obtained at 3, 24, 48, 72, and 96 h post-injection demonstrate radiotracer accumulation at sites corresponding to mCherry-positive tumors (dashed circles); **(C)** Quantitative volume-of-interest (VOI) analysis demonstrates sustained tumor uptake, with increasing radiotracer accumulation at later time point; **(D)** Quantitative VOI analysis of liver-associated signal demonstrates progressive radiotracer clearance over time, resulting in improved tumor-to-liver contrast.

Finally, to characterize the *in vivo* pharmacokinetic profile of the radiotracer, tissue biodistribution studies were performed at 24, 48, and 96 h p.i.. Organ-level analysis demonstrated a distinct and favorable biodistribution pattern for [⁸⁹Zr]Zr-DFO-anti-TRA scFv-Fc across all evaluated tissues. As shown in the biodistribution profile (Fig. 6), the liver exhibited the highest radiotracer uptake, consistent with the hepatic clearance commonly observed for Fc-containing antibody fragments. The spleen showed the second-highest uptake, likely reflecting metastatic tumor involvement, as the spleen is a common site of metastatic implantation in this model ^30,31,33^. Importantly, blood retention at 96 h p.i. was relatively low, indicating efficient systemic clearance of the radiotracer and supporting the generation of high tumor-to-background contrast during PET imaging. Furthermore, most normal tissues including the heart, lungs, kidneys, stomach, pancreas, small intestine, large intestine, bone, and muscle demonstrated minimal nonspecific radiotracer uptake. These results highlight the favorable pharmacokinetic characteristics of the construct and suggest reduced off-target exposure relative to conventional full-length antibody-based imaging agents.

**Figure 6.**
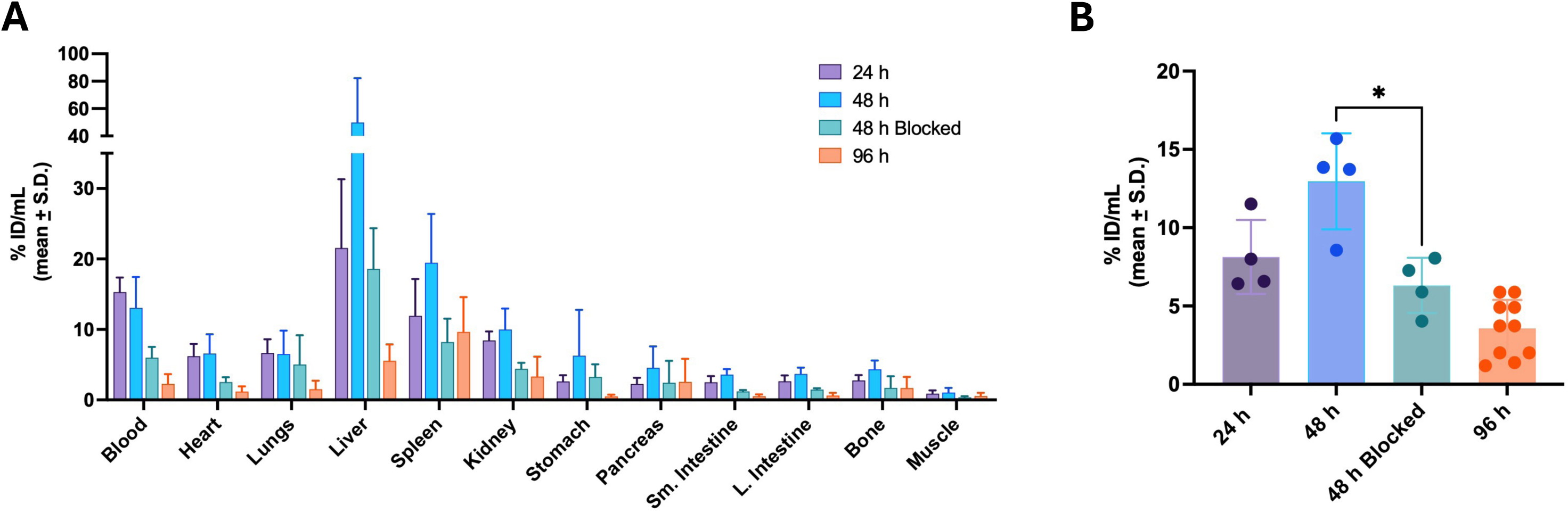
*Ex vivo* biodistribution of [^89^Zr]Zr-DFO-anti-TRA scFv-Fc. **(A)** Mice bearing i.p. TKO tumors received a single i.p. dose (300 μCi) of [^89^Zr]Zr-DFO-anti-TRA scFv-Fc. Tissue uptake (%ID/g) was measured at 24 (n=4), 48 (n=4), and 96 h (n=10) p.i.; **(B)** High uptake of the radiotracer in omental metastasis is observed at 48 h p.i. Competitive binding at 48 h p.i. showed reduced tumor uptake in the group co-administered with the radiotracer and 100-fold excess of unmodified anti-TRA scFv-Fc (n=4), confirming target specificity. * p<0.05.

Collectively, these findings demonstrate that [⁸⁹Zr]Zr-DFO-anti-TRA scFv-Fc achieves selective tumor targeting, sustained tumor retention, and favorable whole-body biodistribution in ovarian cancer models. Importantly, its ability to preferentially accumulate and persist within tumors while clearing from non-target tissues supports its translational potential as an immunoPET agent for noninvasive detection and assessment of TRA-expressing disease. Moreover, these favorable targeting properties establish a strong foundation for adapting the TRA-targeted scFv-Fc platform for radiopharmaceutical therapy, providing a potential theranostic strategy for OC.

## Discussion

The success of molecular targeting depends on the availability of tumor-specific targets that are broadly expressed, readily accessible on the cell surface, and minimally expressed in normal tissues. In this study, we identify the cancer-associated TRA glycoepitope as a promising targeting agent for OC. TRA expression was significantly higher in ovarian tumors than in normal adjacent ovarian tissue and was maintained across histologic subtypes and disease stages, suggesting that its expression is a common feature of epithelial OC rather than being restricted to a particular subtype or stage of disease. In addition, using both human and murine OC models, we further demonstrated that TRA is readily accessible on the tumor cell surface, exhibits efficient antibody penetration throughout 3D tumor spheroids, and supports selective tumor targeting *in vivo*. Finally, radiolabeling of the TRA-targeted scFv-Fc with ^89^Zr generated an immunoPET tracer with sustained tumor retention, favorable pharmacokinetics, and minimal uptake in normal tissues. Collectively, these findings establish TRA as a highly selective and accessible molecular target for OC.

A major strength of TRA as a therapeutic target lies in its recognition of a cancer-associated glycoepitope rather than the underlying protein scaffold. Many cell-surface proteins currently being explored for OC targeting, including folate receptor-α, mesothelin, Trop-2, and podocalyxin, are also expressed in normal tissues, raising concerns regarding off-target toxicity and limiting their therapeutic window. In contrast, TRA recognizes a tumor-specific glycan epitope ^38–40^ generated through aberrant glycosylation, one of the defining hallmarks of malignant transformation ^21,22^. Because abnormal glycosylation is widespread across epithelial malignancies ^21^, targeting cancer-associated glycans offers an opportunity to achieve greater tumor specificity than strategies directed against the protein backbone alone. Our finding that TRA was largely absent from normal adjacent ovarian tissue while remaining broadly expressed across OC histotypes supports this concept and highlights the potential advantage of glycoepitope-directed targeting.

An important finding of this study is that TRA is not only expressed by ovarian tumors but is also highly accessible for antibody-mediated targeting. Successful molecular targeting requires targets that are abundantly expressed on the cell surface and remain accessible within the complex 3D tumor microenvironment. We demonstrated robust penetration of the TRA-targeted scFv-Fc throughout intact tumor spheroids generated from both human and murine OC cells, despite marked differences in spheroid architecture. Furthermore, orthotopic tumors retained strong cell-surface TRA expression, and the Cy7-conjugated anti-TRA scFv-Fc- selectively accumulated within disseminated i.p. tumor implants while exhibiting minimal retention in normal tissues. These findings suggest that TRA remains accessible throughout tumor progression and is suitable for antibody-based imaging and therapeutic applications.

The favorable *in vivo* behavior of the anti-TRA scFv-Fc was further confirmed by PET imaging. Radiolabeling with ^89^Zr yielded a highly stable tracer that demonstrated sustained tumor uptake and prolonged retention while maintaining low background activity in most normal tissues. Importantly, the engineered scFv-Fc format was specifically developed to improve pharmacokinetics by reducing the prolonged circulation associated with full-length IgG antibodies ^38^.The resulting biodistribution profile produced excellent tumor-to-background contrast with low circulation of the scFv-Fc radiotracer in the blood pool at 96 h p.i., supporting the use of this platform for quantitative whole-body imaging. These characteristics are particularly advantageous for OC, which frequently presents with multifocal peritoneal dissemination.

Beyond diagnostic imaging, our findings have important implications for the development of TRA-directed radiopharmaceutical therapy (RPT). The selective and sustained accumulation of the anti-TRA scFv-Fc from the PET images in OC sets the stage forr replacing the diagnostic radionuclide (^89^Zr) with DNA damaging therapeutic radionuclides like ^177^Lutetium (t_1/2_ ∼ 6.67 d). Because TRA expression was maintained across OC histotypes and disease stages while exhibiting minimal expression in normal ovarian tissue, TRA-directed RPT could potentially deliver tumoricidal radiation selectively to disseminated peritoneal disease while minimizing exposure of healthy tissues. Such an approach may be particularly valuable for patients with recurrent OC, where diffuse microscopic peritoneal metastases remain a major therapeutic challenge. Moreover, the same targeting construct could support a true theranostic strategy, in which ^89^Zr immunoPET is used to identify patients with TRA-positive tumors and monitor therapeutic response.

In conclusion, this study identifies the TRA glycoepitope as a highly promising molecular target for OC. More importantly, these findings establish the foundation for a TRA-directed theranostic platform that has the potential to improve both the diagnosis and treatment of OC through highly selective RPT.

## Acknowledgements

This work is supported in part by the Janet Burros Memorial Foundation, the Michigan Ovarian Cancer Alliance, and the Molecular Therapeutics Program at Karmanos Cancer Institute (KCI). The Molecular Therapeutics Program, Biobanking and Correlative Sciences Core, Microscopy, Imaging, and Cytometry Resources Core, and Animal Model and Therapeutics Evaluation Core at KCI are supported in part by NIH Center grant P30 CA022453 to KCI at Wayne State University.

## References

1 Webb, P. M. & Jordan, S. J. Global epidemiology of epithelial ovarian cancer. Nat Rev Clin Oncol 21, 389–400 (2024). 10.1038/s41571-024-00881-3

2 Cabasag, C. J. et al. Ovarian cancer today and tomorrow: A global assessment by world region and Human Development Index using GLOBOCAN 2020. Int J Cancer 151, 1535–1541 (2022). 10.1002/ijc.34002

3 Berek, J. S., Renz, M., Kehoe, S., Kumar, L. & Friedlander, M. Cancer of the ovary, fallopian tube, and peritoneum: 2021 update. Int J Gynaecol Obstet 155 **Suppl 1**, 61–85 (2021). 10.1002/ijgo.13878

4 Markman, M. Pharmaceutical Management of Ovarian Cancer: Current Status. Drugs 79, 1231–1239 (2019). 10.1007/s40265-019-01158-1

5 Armbruster, S., Coleman, R. L. & Rauh-Hain, J. A. Management and Treatment of Recurrent Epithelial Ovarian Cancer. Hematol Oncol Clin North Am 32, 965–982 (2018). 10.1016/j.hoc.2018.07.005

6 DiSilvestro, P. & Alvarez Secord, A. Maintenance treatment of recurrent ovarian cancer: Is it ready for prime time? Cancer Treat Rev 69, 53–65 (2018). 10.1016/j.ctrv.2018.06.001

7 Wang, Z. B. et al. Immunotherapy and the ovarian cancer microenvironment: Exploring potential strategies for enhanced treatment efficacy. Immunology 173, 14–32 (2024). 10.1111/imm.13793

8 Zhang, S. et al. Radiopharmaceuticals and their applications in medicine. Signal Transduct Target Ther 10, 1 (2025). 10.1038/s41392-024-02041-6

9 Varghese, T. P., John, A. & Mathew, J. Revolutionizing cancer treatment: The role of radiopharmaceuticals in modern cancer therapy. Precis Radiat Oncol 8, 145–152 (2024). 10.1002/pro6.1239

10 Miller, S. R., Dewaraja, Y., Rowe, S. P., Reichert, Z. R. & Viglianti, B. L. Review of radiopharmaceutical therapy in prostate cancer. Urologic oncology, 111030 (2026). 10.1016/j.urolonc.2026.111030

11 Wang, J., Pang, X., Lian, J. & Lu, H. (177)Lu-labelled peptide receptor radionuclide therapy in patients with neuroendocrine tumors: a systematic review and meta-analysis. Front Endocrinol (Lausanne) 17, 1758639 (2026). 10.3389/fendo.2026.1758639

12 Crabbe, M., Opsomer, T., Vermeulen, K., Ooms, M. & Segers, C. Targeted radiopharmaceuticals: an underexplored strategy for ovarian cancer. Theranostics 14, 6281–6300 (2024). 10.7150/thno.99782

13 Banerjee, S., Drapkin, R., Richardson, D. L. & Birrer, M. Targeting NaPi2b in ovarian cancer. Cancer Treat Rev 112, 102489 (2023). 10.1016/j.ctrv.2022.102489

14 Hu, X. et al. Advances in the Application of Radionuclide-Labeled HER2 Affibody for the Diagnosis and Treatment of Ovarian Cancer. Front Oncol 12, 917439 (2022). 10.3389/fonc.2022.917439

15 Ritch, S. J. & Telleria, C. M. The Transcoelomic Ecosystem and Epithelial Ovarian Cancer Dissemination. Front Endocrinol (Lausanne) 13, 886533 (2022). 10.3389/fendo.2022.886533

16 Yousefi, M. et al. Current insights into the metastasis of epithelial ovarian cancer - hopes and hurdles. Cell Oncol (Dordr) 43, 515–538 (2020). 10.1007/s13402-020-00513-9

17 AK, M. in Tumor Metastasis (ed Xu K) (Intech, 2016).

18 Hendershot, A. et al. Strategies for prevention and management of ocular events occurring with mirvetuximab soravtansine. Gynecol Oncol Rep 47, 101155 (2023). 10.1016/j.gore.2023.101155

19 Le Tran, N., Wang, Y., Bilandzic, M., Stephens, A. & Nie, G. Podocalyxin promotes the formation of compact and chemoresistant cancer spheroids in high grade serous carcinoma. Sci Rep 14, 7539 (2024). 10.1038/s41598-024-57053-7

20 Le Tran, N., Wang, Y. & Nie, G. Podocalyxin in Normal Tissue and Epithelial Cancer. Cancers 13 (2021). 10.3390/cancers13122863

21 Pinho, S. S. & Reis, C. A. Glycosylation in cancer: mechanisms and clinical implications. Nat Rev Cancer 15, 540–555 (2015). 10.1038/nrc3982

22 Reily, C., Stewart, T. J., Renfrow, M. B. & Novak, J. Glycosylation in health and disease. Nat Rev Nephrol 15, 346–366 (2019). 10.1038/s41581-019-0129-4

23 Gogoi, R. P. et al. A Novel Role of Connective Tissue Growth Factor in the Regulation of the Epithelial Phenotype. Cancers 15 (2023). 10.3390/cancers15194834

24 Chehade, H. et al. MNRR1 is a driver of ovarian cancer progression. Transl Oncol 29, 101623 (2023). 10.1016/j.tranon.2023.101623

25 Tedja, R. et al. Protein kinase Cα-mediated phosphorylation of Twist1 at Ser-144 prevents Twist1 ubiquitination and stabilizes it. J Biol Chem 294, 5082–5093 (2019). 10.1074/jbc.RA118.005921

26 Cardenas, C. et al. Adipocyte microenvironment promotes Bclxl expression and confers chemoresistance in ovarian cancer cells. Apoptosis 22, 558–569 (2017). 10.1007/s10495-016-1339-x

27 Alvero, A. B. et al. TRX-E-002-1 Induces c-Jun-Dependent Apoptosis in Ovarian Cancer Stem Cells and Prevents Recurrence In Vivo. Mol Cancer Ther 15, 1279–1290 (2016). 10.1158/1535-7163.MCT-16-0005

28 Kim, J., Coffey, D. M., Ma, L. & Matzuk, M. M. The ovary is an alternative site of origin for high-grade serous ovarian cancer in mice. Endocrinology 156, 1975–1981 (2015). 10.1210/en.2014-1977

29 Craveiro, V. et al. Phenotypic modifications in ovarian cancer stem cells following Paclitaxel treatment. Cancer Med 2, 751–762 (2013). 10.1002/cam4.115

30 Fox A, L. G., Adzibolosu N, Wong T, Tedja R, Sharma S, Gogoi R, Morris R, Mor G, Fehl C, Alvero AB. Adipose microenvironment promotes hypersialylation of ovarian cancer cells. Frontiers in Oncology 14 (2024). 10.3389/fonc.2024.1432333

31 Alvero, A. B. et al. Immune Modulation of Innate and Adaptive Responses Restores Immune Surveillance and Establishes Antitumor Immunologic Memory. Cancer Immunol Res 12, 261–274 (2024). 10.1158/2326-6066.CIR-23-0127

32 van den Pol, A. N., et al. Lassa-VSV chimeric virus targets and destroys human and mouse ovarian cancer by direct oncolytic action and by initiating an anti-tumor response. Virology 555, 44–55 (2021). 10.1016/j.virol.2020.10.009

33 Alvero, A. B. et al. Transimmunization restores immune surveillance and prevents recurrence in a syngeneic mouse model of ovarian cancer. Oncoimmunology 9, 1758869 (2020). 10.1080/2162402x.2020.1758869

34. (!!! INVALID CITATION !!! 30).

35 Tateno, H. et al. Podocalyxin is a glycoprotein ligand of the human pluripotent stem cell-specific probe rBC2LCN. Stem Cells Transl Med 2, 265–273 (2013). 10.5966/sctm.2012-0154

36 Hernandez, L. et al. Characterization of ovarian cancer cell lines as in vivo models for preclinical studies. Gynecol Oncol 142, 332–340 (2016). 10.1016/j.ygyno.2016.05.028

37 Kim, J. et al. High-grade serous ovarian cancer arises from fallopian tube in a mouse model. Proc Natl Acad Sci U S A 109, 3921–3926 (2012). 10.1073/pnas.1117135109

38 White, J. M. et al. Selective ablation of TRA-1-60(+) pluripotent stem cells suppresses tumor growth of prostate cancer. Theranostics 13, 2057–2071 (2023). 10.7150/thno.78915

39 White, J. M. et al. Detecting TRA-1-60 in Cancer via a Novel Zr-89 Labeled ImmunoPET Imaging Agent. Mol Pharm 17, 1139–1147 (2020). 10.1021/acs.molpharmaceut.9b01181

40 Schopperle, W. M. & DeWolf, W. C. The TRA-1-60 and TRA-1-81 human pluripotent stem cell markers are expressed on podocalyxin in embryonal carcinoma. Stem Cells 25, 723–730 (2007). 10.1634/stemcells.2005-0597

